# The p97 adaptor p47/NSFL1C is necessary for stress granule dissolution after heat stress

**DOI:** 10.64898/2026.06.08.730917

**Authors:** Michelle A. Johnson, Richa Khanna, Sirisha Mukkavalli, Ly Nguyen, Malavika Raman

## Abstract

Stress granules form in response to diverse cellular perturbations to sequester translation components until the stress is resolved. Stress granules are composed of RNA-protein assemblies in membrane delimited structures and must be rapidly disassembled to release components to allow translation to resume. Disassembly of stress granules formed in response to heat stress is dependent on ubiquitiylation of stress granule components such as G3BP1. Ubiquitylation of stress granule proteins recruits the AAA-ATPase p97 (also known as VCP) to enable ubiquitin-dependent disassembly of these structures. Loss of p97 activity leads to the persistence of stress granules and is implicated in several age-related neurodegenerative diseases. Here we show that p97 recruitment to stress granules is dependent on its ubiquitin binding co-factor p47. p47 translocates to stress granules in response to a variety of cellular stressors and is required for the recruitment of p97 to stress granules. Loss of p47 leads to an inhibition in stress granule disassembly. We further show that p47 associates with G3BP1 in response to heat stress in a ubiquitin-dependent manner. Taken together our data adds to the growing list of p97 adaptors that are implicated in the recruitment of p97 for dissolution of stress granules.

## Introduction

Translation is an energy-intensive process that is finely tuned to cellular stress. When cells experience adverse environments, for example viral infection, oxidative or heat stress, they temporarily stall translation to conserve energy until the stress is resolved (Hershey, Sonenberg, & Mathews, 2019). Translation attenuation occurs through multiple overlapping processes, including ribosome splitting and inhibition of eIF2 mediated pre-initiation complex formation. Another mechanism is to sequester mRNAs, RNA binding proteins and ribosome subunits into a membrane-less phase separated structure in the cytoplasm known as the stress granule (Banani, Lee, Hyman, & Rosen, 2017; Bharath et al., 2020; Anthony A. Hyman, Christoph A. Weber, & Frank Jülicher, 2014). These structures rapidly form in response to cellular stress but are efficiently dissolved to allow cells to rapidly resume translation when the stress is resolved.

Stress granule formation depends on a core set of proteins that bind RNA. These RNA binding proteins (such as G3BP1/2 or TIA1) contain intrinsically disordered regions (IDRs) that self-associate and allow for protein and RNA to coalesce through a process known as liquid-liquid phase separation (LLPS) and form a droplet-like structure in the cytoplasm (Anthony A. Hyman et al., 2014; T. Mittag & Parker, 2018). LLPS is facilitated by protein-protein, protein-RNA and RNA-RNA interactions. Studies report that the IDR interactions must cross what is known as a percolation threshold to enable LLPS (Tanja Mittag & Pappu, 2022; Pappu, Cohen, Dar, Farag, & Kar, 2023). Stress granules can vary in size, composition and structure and usually contain on the order of hundreds of resident proteins (Hu et al., 2023; Jain et al., 2016; Markmiller et al., 2018; Marmor-Kollet et al., 2020). The core of the stress granule is quite stable and composed of group of RNA binding proteins with IDRs that is surrounded by a more dynamic shell of loosely associated proteins (Banani et al., 2017). There are about seven core stress granule proteins routinely identified from loss of function screens and proteomics. Interestingly loss of any one of these is not necessary for stress granule formation, indicating a degree of redundancy. However, key stress granule organizers are G3BP1 and 2 (Kedersha et al., 2016; Yang et al., 2020). While loss of either does not impact stress granule formation, a double knock out has been shown to completely abolish stress granule formation.

Given the transient nature of stress granules and the important proteins and RNAs they sequester, it is paramount that these structures dissolve once the stressor is removed. It is now appreciated that when stress granules are not appropriately resolved, they can form pathological entities that are harmful for the cell. Such aberrant stress granules form in the context of diverse disease states from cancer to neurodegeneration (Mateju et al., 2017; Patel et al., 2015). Furthermore, proteins that normally participate in LLPS are mutated in disease and importantly, the disease-associated variants promote aberrant phase transitions. This is a prominent feature in neurodegenerative diseases where mutations in α-synuclein, huntingtin, hnRNPA1, TDP43 and FUS can lower the threshold necessary for LLPS (Zbinden, Pérez-Berlanga, De Rossi, & Polymenidou, 2020). In pathological LLPS, the phase separated structures lose their liquid-like property and are unable to dissipate. Over time, these structures transition into a more solid-like state amyloid that is detrimental to cell function and viability.

While we know what factors contribute to stress granule assembly, less is known about how these structures are resolved. Recent work suggests that the context in which stress granules form can lead to post-translational modifications of proteins within these structures that influences how they are dissipated. Stress granules are ubiquitylated to differing degrees depending on the type of stress, with heat stress inducing the most robust ubiquitylation of these structures (Gwon et al., 2021; Maxwell et al., 2021; Tolay & Buchberger, 2021). Studies have found that stress granule proteins can be decorated with free monoubiquitin, unanchored ubiquitin chains, as well proteins covalently modified with K48 and K63 linked ubiquitin chains (Gwon et al., 2021; Markmiller et al., 2019; Maxwell et al., 2021; Xie et al., 2018). Notably, heat stress, but not oxidative stress causes robust ubiquitylation of G3BP1 and other stress granule proteins via K63-linked ubiquitin chains (Gwon et al., 2021; Maxwell et al., 2021). Ubiquitylated of G3BP1 serves as the signal for the recruitment of factors that aid in stress granule disassembly (Gwon et al., 2021). A key component of ubiquitin-mediated stress granule disassembly is the p97 AAA-ATPase (also known as valosin containing protein, VCP or Cdc48p in yeast) (Buchan, Kolaitis, Taylor, & Parker, 2013). p97 is an abundant homo-hexameric ATPase that maintains protein homeostasis (proteostasis) within the cell (Ahlstedt, Ganji, & Raman, 2022; H. Meyer, 2012). p97 recognizes ubiquitylated proteins via a host of dedicated adaptors (estimated to be over 30 in mammalian cells) (Buchberger, Schindelin, & Hanzelmann, 2015). These adaptors serve as bridges that recruit p97 via their dedicated interaction domains such as ubiquitin X domain (UBX) or short linear motifs to ubiquitylated substrates that they associate with via ubiquitin associated domains (UBA). Consecutive ATP hydrolysis via p97 leads to the unfolding of the proximal ubiquitin on the substrate that proceeds in the direction of the substrate such that it is pulled through the central pore and unfolded (Cooney et al., 2019; Twomey et al., 2019). This activity is a prerequisite for degradation of many substrates by the 26S proteasome. The prominent p97 adaptor dimer UFD1-NPL4 is required for orienting the ubiquitin chain on the substrate to begin unfolding (Braxton & Southworth, 2023). UFD1-NPL4 has been shown to be necessary for stress granule disassembly (Buchan et al., 2013). However, it is not the only p97 adapter involved in stress granule disassembly, as recently another adaptor UBXD8 (also known as FAF2) was also found to recruit p97 to ubiquitylated G3BP1 to initiate disassembly (Gwon et al., 2021). UBXD8 is an ER embedded adaptor that primarily recruits p97 for ER-associated degradation which is the clearance of misfolded proteins from the ER (Ganji et al., 2023; Loregger et al., 2017; Nahar, Chowdhury, Ogura, & Esaki, 2020; Xu, Liu, Lee, & Ye, 2013). Intriguingly, the ER has been demonstrated to tether stress granules; whether this is via UBXD8 interaction with G3BP1 is presently unknown (Lee, Cathey, Wu, Parker, & Voeltz, 2020).

In this study we find that another p97 adaptor known as p47 (or NSFL1C) is also recruited to stress granules and required for their disassembly. In an imaging screen for p97 adaptors that contain UBA-UBX domains we found several adaptors were recruited to stress granules induced by heat shock and oxidative stress including UBXD1, UBXD8 and p47. Loss of p47 decreased p97 recruitment to heat shock induced stress granules and as a result interrupted stress granule dissolution after stress removal. We further show that p47 interacts with G3BP1 and is recruited to stress granules in a manner dependent on ubiquitylation. Taken together, our results suggest that multiple adaptors recruit p97 in a redundant manner to stress granules ensuring efficient resolution of these structures when stress is resolved.

## Results

### p97 is recruited to stress granules induced by heat shock and oxidative stress

p97 and its yeast homolog Cdc48p have been demonstrated to be essential for the dissolution of stress granules (Buchan et al., 2013; Gwon et al., 2021; Wang et al., 2019). To investigate the role of p97 adapters in stress granule assembly and disassembly, we engineered HeLa Kyoto cells to stably expressed low levels of GFP-G3BP1 with lentivirus such that treatment with cellular stress resulted in the rapid formation of phase separated stress granules. GFP-G3BP1 Hela cells were treated with either heat shock (42 °C, 1 hour) or oxidative stress (500 μM sodium arsenite 1 hour) and cells were fixed and stained for p97 (Supplementary Figure 1A). The colocalization of p97 with GFP-G3BP1 was determined by Pearson’s correlation coefficient. In untreated cells p97 was largely cytosolic or ER localized, but in heat shock or sodium arsenite treated cells, there was a re-localization of p97 to punctate, GFP-G3BP1 positive structures (Supplementary Figure 1A and B). Recent studies report that heat shock induces robust ubiquitylation of stress granule proteins (primarily G3BP1 and 2) whereas oxidative stress does not. Furthermore, ubiquitylation is required for the disassembly of stress granules after stress dissipation through a p97-dependent manner. We sought to confirm these findings in Hela and U2OS cells treated with either heat shock or sodium arsenite that were then fixed and stained with ubiquitin and G3BP1. Indeed, while heat shock resulted in increased ubiquitin co-staining with G3BP1 positive foci there was no ubiquitin observed to co-localize with G3BP1-positive stress granules in sodium arsenite treated cells (Supplementary Figure 1C).

To verify the catalytic role of p97 in this process we focused on heat shock induced stress granules as they are known to be ubiquitylated (Gwon et al., 2021; Maxwell et al., 2021). Hela GFP-G3BP1 cells were pre-treated inhibitors to either the proteasome (Bortezomib, Btz, 5μM), p97 (CB5083, 10 μM) or the ubiquitin E1 enzyme (TAK243, 2μM), followed by heat shock. Cells were either fixed or further released at 37 °C for 5 hours for stress granules to dissipate in the presence of absence of the inhibitors (Supplementary Figure 1D, top panel). We found that stress granule formation was not perturbed in the presence of any of the inhibitors in agreement with previous findings (Gwon et al., 2021; Maxwell et al., 2021; Tolay & Buchberger, 2021) (Supplementary Figure 1D, red bars). While stress granules cleared within 5 hours of return to 37 °C, the presence of the inhibitors during the release period prevented recovery and stress granules persisted (Supplementary Figure 1D, green bars). Taken together, ubiquitylation of stress granules, p97 catalytic activity and proteasomal degradation are required for the normal clearance of heat shock induced stress granules.

### Multiple p97 adaptors including p47 and UBXD1 are recruited to stress granules

We next asked which p97 adaptors were recruited to stress granules. The UFD1-NPL4 dimer is known to be required for stress granule dissolution with conserved functions from yeast to mammalian cells (Buchan et al., 2013). As expected NPL4 colocalized to both heat shock and sodium arsenite induced stress granules (Figure 1 A, B). We were unable to test UFD1 localization due to the lack of antibodies that worked well for immunofluorescence. As previously reported, UBXD8 also robustly localized to stress granules in response to both stressors (Figure 1C and D) (Gwon et al., 2021). Surprisingly, we found that UBXD1 and p47, two p97 adaptors not previously known to reside at stress granules were recruited to these structures in response to both heat shock and sodium arsenite (Figure 1E-H). UBXD1 is best understood in recruiting p97 to lysosomes to initiate selective degradation of damaged lysosomes by autophagy (lysophagy) (Kirchner, Bug, & Meyer, 2013; Klickstein et al., 2024; Papadopoulos et al., 2017) and p47 is the p97 adaptor that regulates the reformation of the Golgi apparatus after mitosis (Kondo et al., 1997; H. H. Meyer, Kondo, & Warren, 1998; Uchiyama & Kondo, 2005). To ensure that UBXD1 and p47 adaptor recruitment was specific, we additionally stained for UBXD7 and found that it was not localized to stress granules with either stressor (Figure 1I and J). We additionally verified that UBXD1 and p47 recruitment was not unique to HeLa Kyoto cells. UBXD1 and p47 were recruited to stress granules in mouse embryonic fibroblasts, U2OS cells treated with heat shock and sodium arsenite (Supplementary Figure 2A-D).

**Figure 1.**
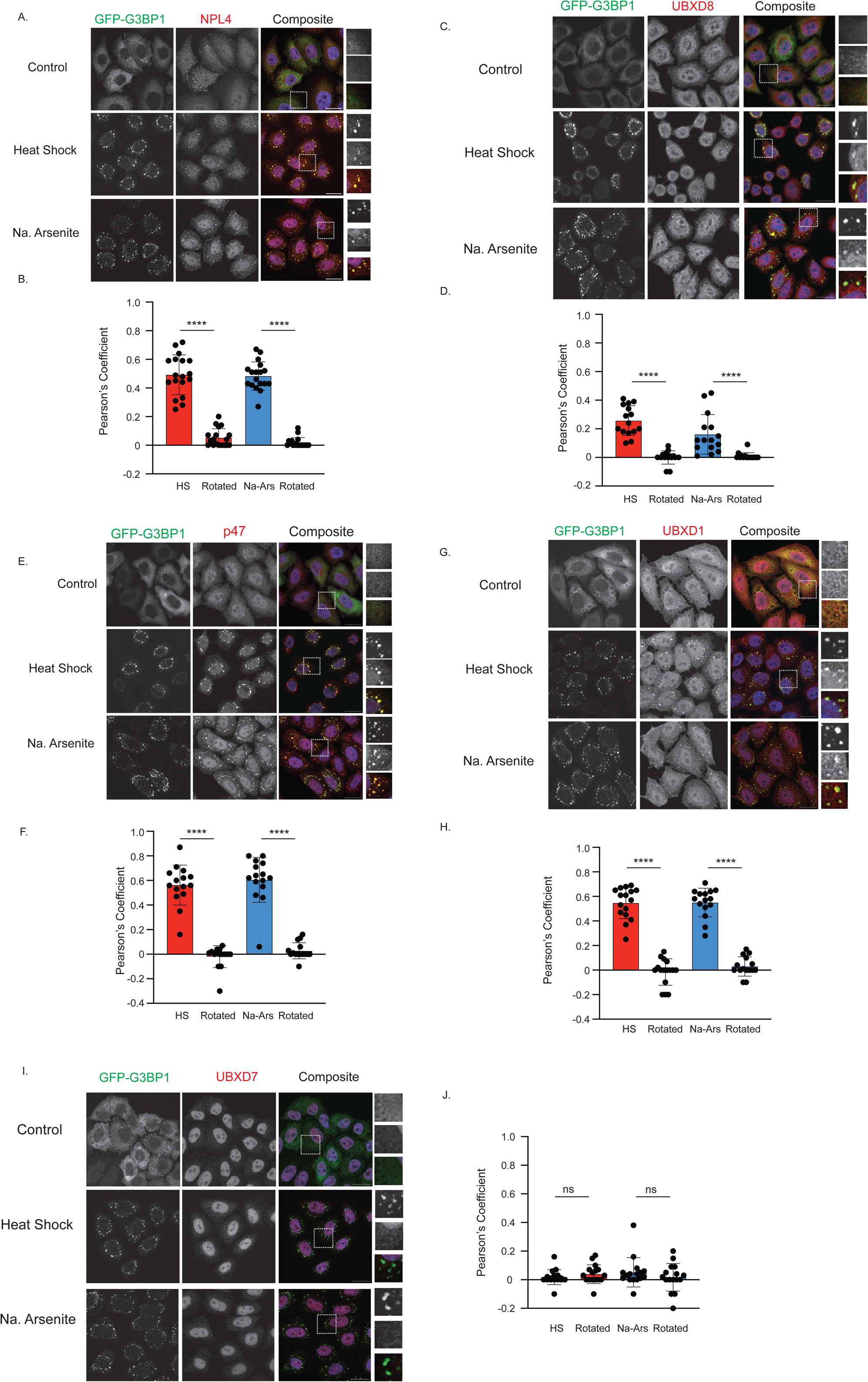
Multiple p97 adaptors including p47 and UBXD1 are recruited to stress granules induced by heat shock and sodium arsenite. Recruitment of p97 adapters to GFP-G3BP1 positive punctate in HeLa Kyoto cells was visualized via immunofluorescent staining of (A) NPL4 (C) UBXD8 (E) p47 (G) UBXD1 and (I) UBXD7 following Heat Shock (HS, 42°C for 1 hour) or sodium arsenite (Na Ars., 500μM for 1 hour). Nuclei were counterstained with Hoechst (B, D, F, H, J). Colocalization is numerically represented via the Pearson’s correlation coefficient (PCC) of G3BP1 positive punctate with: (B) NPL4 (D) UBXD8 (F) p47 (H) UBXD1 (J) UBXD7. The significance of colocalization was determined by comparing the PCC for unaltered images to the PCC of a matched but rotated image, termed “Rotated”. N= 3 biological replicates where 5 images were quantified per condition and is represented as mean PCC +/- Standard Deviation. Significance was determined via a paired t-test, *: p<0.05, ns: not significant. Scale bar: 20μm.

### p47 but not UBXD1 is required for stress granule disassembly following heat shock

The recruitment of p47 and UBXD1 to stress granules prompted us to investigate if these were bystanders or were required for formation or dissolution of these structures. To test this, we depleted p47, UBXD1 or UBXD8 using siRNAs in GFP-G3BP1 cells (Figure 2A and Supplementary Figure 3). UBXD8 was used as a positive control as it is already known to be necessary for stress granule disassembly. Cells were treated with heat shock (1 hour) and released (5 hours) and then fixed and imaged. The number of cells with GFP-G3BP1 puncta were quantified. We found that knockdown of p47 or UBXD1 did not perturb stress granule assembly (Figure 2 B and C and Supplementary Figure 3 A and B). However, knockdown of p47 with two independent siRNAs caused an inhibition in the dissolution of stress granules after heat shock relief (Figure 2 B and C). This inhibition of release was comparable to that seen with depletion of UBXD8 (Figure 2 B and C). Surprisingly, even though UBXD1 was robustly recruited to stress granules, it did not appear to be required for their dissolution as its depletion did not perturb stress granule release (Supplementary Figure 3 A and B). We co-depleted p47 and UBXD8 and found no greater inhibition in stress granule release suggesting that these proteins likely do not act in parallel pathways (Figure 2 A-C). To ensure that the inhibition of stress granule release in p47--depleted cells was specific, we next performed rescue experiments. We complemented p47 siRNA transfected cells with a FLAG-tagged p47 cDNA construct that was resistant to the p47 siRNA (Figure 2 D and E). In the p47-rescue transfected cells, release from heat shock resulted in the dissipation of stress granules in a manner comparable to wildtype cells (Figure 2 D-F).

**Figure 2.**
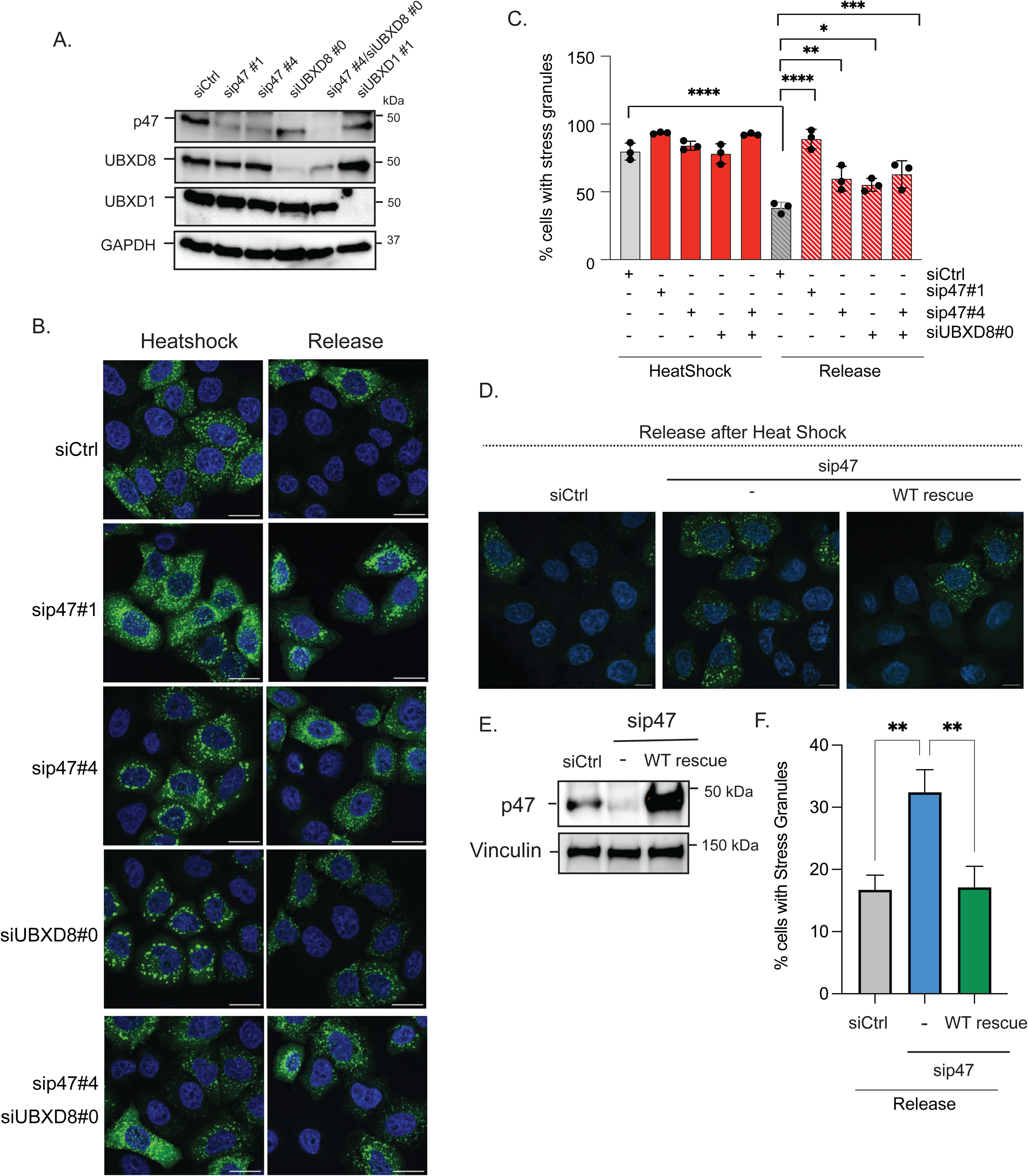
p47 is required for the dissolution of stress granules after heat stress. (A) HeLa Kyoto GFP-G3BP1 cells were transiently transfected with siRNAs to p47 (two siRNAs), UBXD8, UBXD1 or a non-targeting control (Ctrl) for 48 hours. One set of cells was co-depleted of p47 and UBXD8 as indicated. Immunoblot of knockdown efficiency is shown. B. HeLa Kyoto GFP-G3BP1 depleted with siRNAs as indicated in (A) were treated with Heat Shock (42°C for 1 hour) and then released into 37°C for 5 hours. Cells were fixed and imaged for GFP-G3BP1. Nuclei were counterstained with Hoechst. C. Quantification of number of cells with stress granules from (B). (D) HeLa Kyoto GFP-G3BP1 cells were transiently transfected with siRNAs to a non-targeting control (Ctrl) or p47 for 24 hours. p47-depleted cells were transfected with a siRNA resistant p47 cDNA for a further 24 hours. Cells were treated with Heat Shock (42°C for 1 hour) and then released into 37°C for 5 hours. Released cells were fixed and imaged for GFP-G3BP1. Nuclei were counterstained with Hoechst. (E) Immunoblot of p47 levels from (D). (F) Quantification of percentage of cells with stress granules under release conditions from (D). 100-250 cells (panel C) and 100-150 cells (panel F) were analyzed in N=3 independent experiments. Bar graph shows mean and standard error of the mean. *,**,***,****: p< 0.05,0.01,0.001,0.0001 as determined by One way ANOVA with Dunnetts (panel C) or Šidáks (panel F) multiple Comparisons test. Scale bar is 10 μM.

### p47 is required for recruitment of p97 to stress granules

Adaptors are in many instances known to recruit p97 to substrates that require degradation. To determine whether p47 was indeed a recruitment factor for p97 to stress granules, we depleted p47 and evaluated the recruitment of p97 to heat shock induced stress granules. HeLa GFP-G3BP1 cells were transfected with control or p47 siRNAs and then treated with heat shock. We also depleted UBXD8 as it has previously been demonstrated to recruit p97 to stress granules (Gwon et al., 2021). Cells were fixed and stained for p97 and the co-localization of p97 to GFP-G3BP1 stress granules was determined using the Pearsons’s correlation coefficient. As seen in Figure 3A, heat shock resulted in p97 redistribution to GFP-G3BP1 positive foci. However, depletion of p47 but not UBXD8 resulted in decreased p97 recruitment to GFP-G3BP1 foci (Figure 3 A-C). Taken together, p47 is necessary for the recruitment of p97 to heat shock induced stress granules.

**Figure 3.**
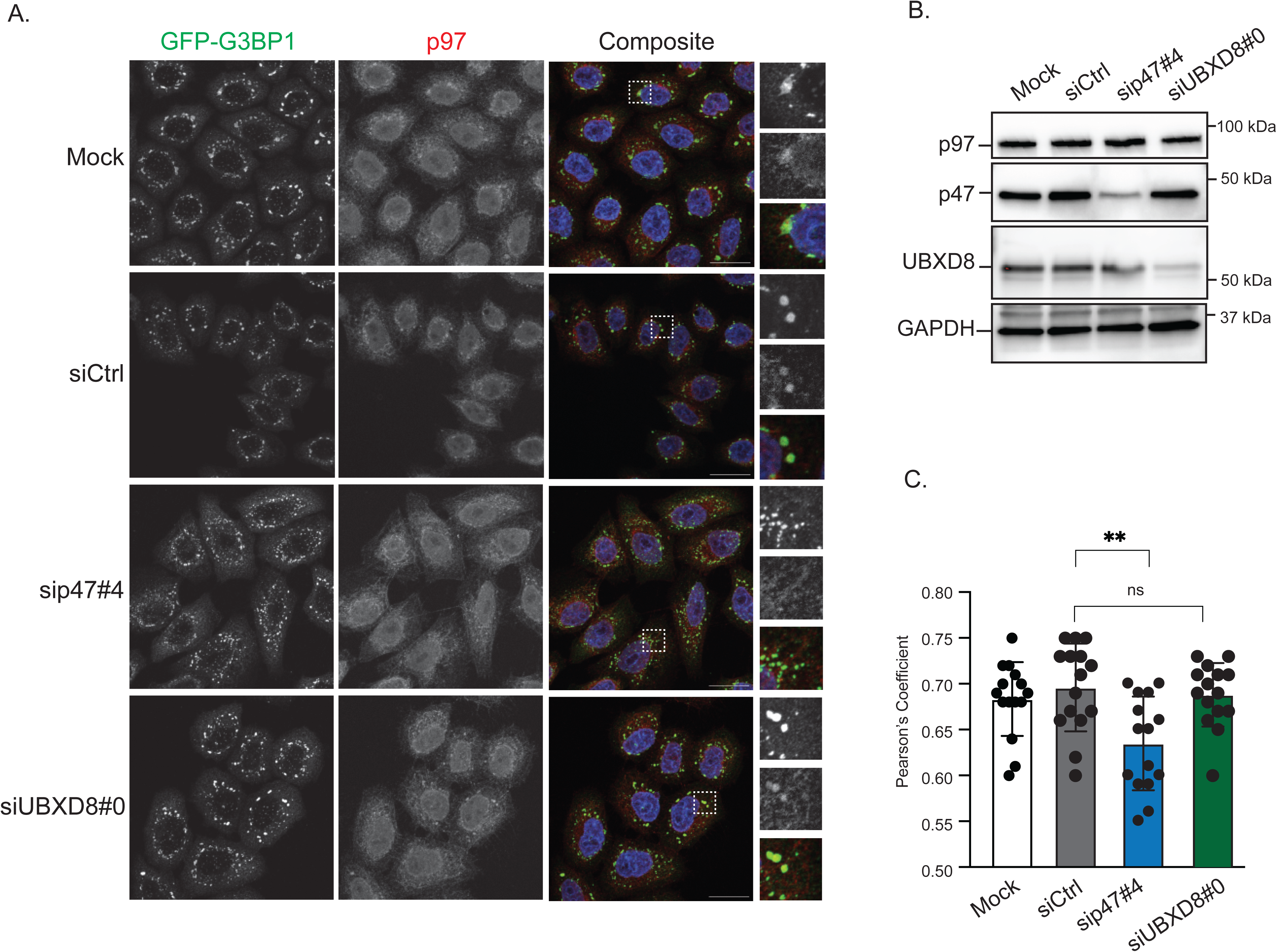
p47 is required for the recruitment of p97 to heat shock induced stress granules. (A) HeLa Kyoto GFP-G3BP1 cells were mock transfected or transfected with non-targeting siRNA (Ctrl), or siRNAs against p47 or UBXD8. Cells were treated with heat shock (42°C for 1 hour) and stained for p97. Depletion of p47 showed disrupted p97 recruitment to G3BP1 positive punctate following heat shock more so than UBXD8 depletion. Nuclei were countered stained with Hoechst. (B) Representative immunoblot of p47 and UBXD8 siRNA depletion. (C) Colocalization represented via Pearson’s Correlation coefficient (PCC) shows reduced p97 recruitment G3BP1 positive punctate following knockdown of p47. Each dot represents an image of a field of at least 5 cells analyzed (for a total of ∼ 70 cells per condition). N= three independent experiments and is shown as mean +/- Standard Deviation. Significance was determined via an Ordinary One-way ANOVA with Tukey’s post hoc test, *: p<0.05, ns: not significant. Scale bar: 20μm

### p47 interacts with G3BP1 in response to heat shock

G3BP1 and 2 are abundant RNA binding proteins and key stress granule organizers. Ubiquitylation of G3BP1 in response to heat shock recruits the p97-UBXD8 complex to initiate the degradation of G3BP1 and disassembly of stress granule. We asked if p47 also interacted G3BP1 in response to heat shock. HeLa cells were treated with either heat shock or sodium arsenite and released (to 37 °C or drug free media respectively) and endogenous G3BP1 was immunoprecipitated from cell lysates. We were able to readily detect p97, UBXD8 and p47 in G3BP1 immunopurifications in a manner stimulated by heat shock. Very little interaction was recovered in control IgG or untreated samples as well as in cells treated with sodium arsenite (Figure 4 A). The interaction between the p97-adaptor complexes and G3BP1 was specific to stress as it was decreased in cells where the stress was resolved (compare lanes 4 and 2 in Figure 4 A). We also performed reciprocal immunopurifications where we purified p47 from an identical set of treated samples and probed for G3BP1. Once again, we were able to detect a heat shock (but not sodium arsenite) induced interaction between p47 and G3BP1. The interaction was weaker under release conditions (Figure 4 B). p47 interacted with p97 under all treatment conditions likely due to other pathways that the two participate in together. Next, we asked whether the interaction between p47 and G3BP1 was dependent of ubiquitin interaction. We pretreated HeLa cells with the ubiquitin E1 inhibitor TAK243 to globally inhibit cellular ubiquitylation prior to treating cells with heat shock. p47 was immunoprecipitated from cells and probed for associated G3BP1. While heat shock induced a strong interaction between p47 and G3BP1, inhibition of ubiquitylation with TAK243 prevented the interaction. To extend this finding further, we asked if the recruitment of p47 to stress granules was dependent on ubiquitination. We treated Hela cells with TAK243 along with heat shock and stained cells for stress granules (G3BP1) and p47. The intensity of p47 at G3BP1 positive foci was determined. Supporting the ubiquitin dependent association between p47 and G3BP1, we found that TAK243 treatment during heat shock decreased the amount of p47 recruited to stress granules (Figure 4 D and E). Taken together, our results suggest that p47 interacts with G3BP1 in a heat shock and ubiquitylation dependent manner.

**Figure 4.**
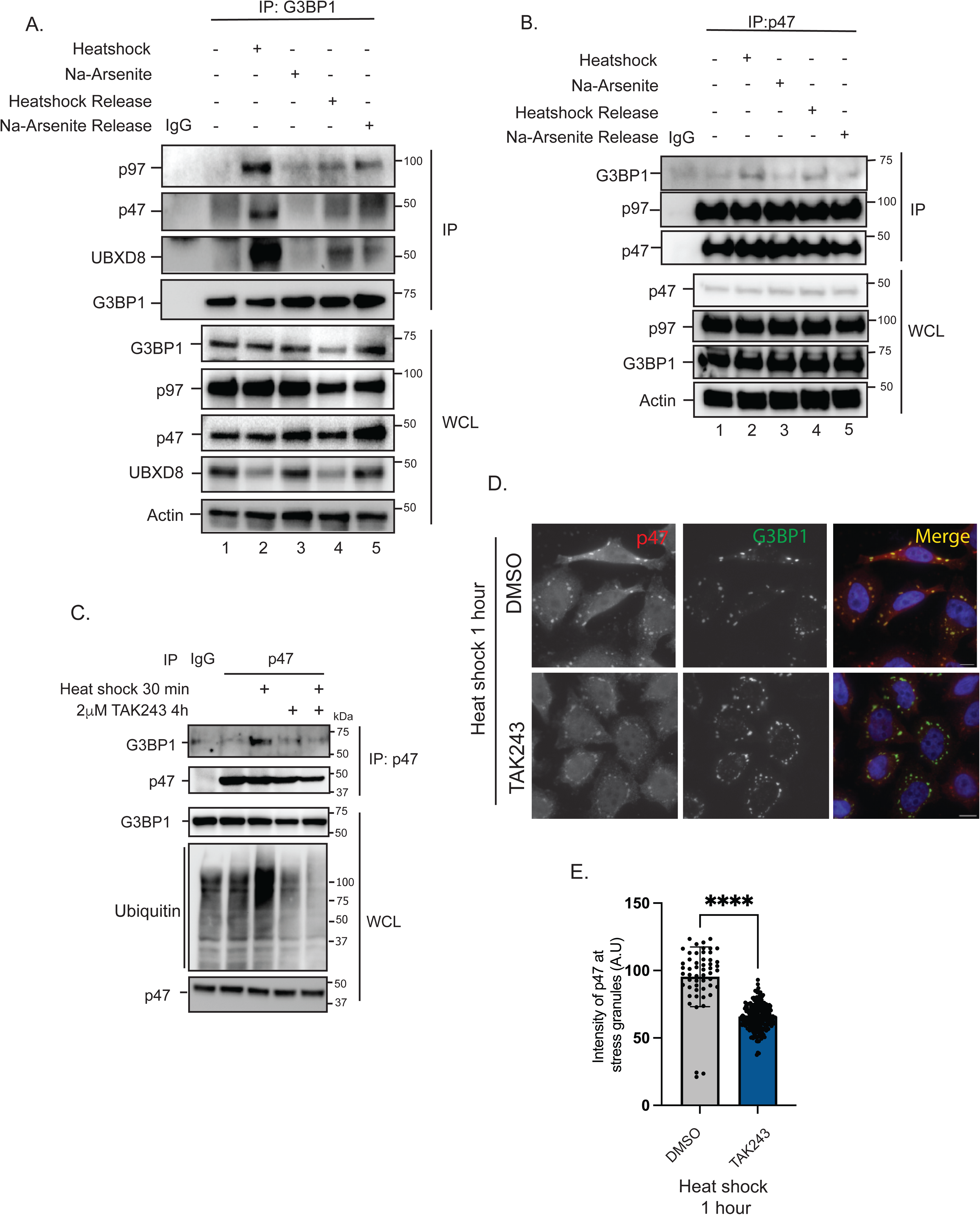
p47 interacts with G3BP1 in a ubiquitin-dependent manner. (A) HeLa Kyoto cells were treated with either Heat Shock (42 °C for 1 hour) and the released (37 °C for 5 hours) or with sodium arsenite (500 μM for 1 hour) and then released into drug free media as indicated. Cells were lysed and endogenous G3BP1 was immunoprecipitated (rabbit IgG was used as a control). Immunoprecipitates were probed for the indicated antibodies. p97, p47 and UBXD8 interact with G3BP1 in a heat shock dependent manner. (B) Similar experiment as in (A) except endogenous p47 was immunoprecipitated. (C) HeLa Kyoto cells were transfected with FLAG-p47 cDNA for 24h. Cells were pre-treated with the ubiquitin E1 inhibitor TAK243 for 4h followed by heat shock (42 °C for 30 min) in the presence of TAK243. Cells were lysed and FLAG-p47 was immunoprecipitated. Immunoprecipitates were probed for G3BP1. G3BP1 interacts with p47 in a heat shock and ubiquitin dependent manner. (D) HeLa Kyoto GFP-G3BP1 cells were pre-treated with the ubiquitin E1 inhibitor TAK243 for 4h followed by heat shock (42 °C for 1 hour) in the presence of TAK243. Cells were fixed and stained for endogenous p47. Nuclei were counterstained with Hoechst. (E) The intensity of p47 at G3BP1 positive stress granules in quantified. Loss of ubiquitination of stress granules diminishes p47 recruitment to stress granules. ∼150 stress granules from 50 cells were analyzed in N=3 independent experiments. Bar graph shows mean and standard deviation. **** : p< 0.0001 as determined by unpaired t test. Scale bar is 10 μM.

## Discussion

The formation and resolution of stress granules is a dynamic process that must be strictly regulated to ensure that translation is limited during times of stress but can rapidly resume when stress is alleviated (Banani et al., 2017; A. A. Hyman, C. A. Weber, & F. Jülicher, 2014). The formation of these dynamic membrane delimited structures is aided by liquid-liquid phase separation and assisted by interactions between low complexity IDRs in RNA binding proteins (T. Mittag & Parker, 2018). While these interactions facilitate the rapid formation of these structures, they can pose a problem to their resolution as evidenced by the formation of aberrant or toxic stress granules in age-associated neurodegenerative disorders (Zbinden et al., 2020). Thus, stress granule disassembly is not a passive process, and requires the concerted action of several proteins, notably the p97 ATPase. Mutations in p97 cause a pleiotropic disorder known as Multisystem Proteinopathy 1 (MSP1) (Johnson, Klickstein, Khanna, Gou, & Raman, 2022; Kimonis, Fulchiero, Vesa, & Watts, 2008). Notably, p97 mutations that cause MSP1 are unable to dissipate stress granules which persist and become toxic over time (Buchan et al., 2013). Furthermore, several proteins that reside in stress granules are known to be mutated in patients with related MSP syndromes or other disorders such as amyotrophic lateral sclerosis, frontotemporal dementia and inclusion body myopathy (IBM). In these disorders, mutant proteins dramatically alter the dynamics of stress granule assembly and disassembly, skewing the rate of disassembly such that these entities persist longer in cells, eventually forming amyloid like fibrils (Custer, Neumann, Lu, Wright, & Taylor, 2010; Kurashige et al., 2021; Nalbandian et al., 2012). These and other studies highlight the interplay between of aberrant phase transitions and protein homeostasis in age associated degenerative diseases.

Intriguingly, recent studies have found that stress granules that arise from different types of perturbations maybe intrinsically distinct (Maxwell et al., 2021). For instance, stress granules formed as a result of heat stress are contain proteins that are modified by ubiquitin whereas when stress granules are formed as a result of sodium arsenite treatment they are largely ubiquitin negative (Gwon et al., 2021; Maxwell et al., 2021). While the net result of stress and granule formation is ostensibly the same, i.e. translation shutoff, it is unclear why different stressors result in these distinct post-translational modifications. The activity of p97 is needed for the resolution of ubiquitin modified stress granules (Gwon et al., 2021). Previous studies have identified the UFD1-NPL4 dimer and UBXD8/FAF2 as being important for the recruitment of p97 to stress granules (Gwon et al., 2021; Liu et al., 2020). In this study we find p47 is an additional p97 adaptor recruited to stress granules that also aids in disassembly of these structures after heat stress.

p47 was originally found in a landmark study several decades ago to regulate Golgi cisternae regrowth from mitotic fragments mitosis (Kondo et al., 1997; H. H. Meyer et al., 1998; Uchiyama & Kondo, 2005). It was found to be the first p97 cofactor for membrane fusion events and a key component of membrane dynamics during cell division. p47 contains a <u>ub</u>iquitin <u>a</u>ssociated (UBA) domain that interacts with ubiquitin on clients, followed by a <u>S</u>hp1, <u>e</u>yes closed (eyc) and <u>p</u>47 (SEP) domain and then the <u>ub</u>iquitin X (UBX) domain for p97 interaction. The SEP domain mediates the self-association of p47 and is responsible for the trimerization of p47 (Yuan et al., 2001). Three p47 molecules bind to a single p97 hexamer and trimerization via the SEP domain is important for the formation of the p47-p97 complex (Beuron et al., 2006; Kondo, Uchiyama, & Okiyoneda, 2004). Notably, upon p97 interaction p47 self-association via the SEP domain is disrupted (Raseekan, Black, & Huang, 2025). The linkers connecting the UBA, SEP and UBX domain are known to be intrinsically disordered (Raseekan et al., 2025). Given that proteins with IDRs have a propensity to phase separate it is tempting to speculate whether one of these linkers may be important for the recruitment of p47 to stress granules.

Collectively our study along that several others have identified a number of p97 adaptors that are recruited to stress granules. This includes UFD1-NPL4, UBXD8/FAF2, ZFAND1 (Turakhiya et al., 2018) and now p47. While we also identified UBXD1 recruitment to stress granules, its depletion did not have any impact on formation or resolution of these structures. A recent study from the Kundu group found that the p97 adaptor ASPL/TUG was required for the assembly of stress granules (Pareek et al., 2025). Why would stress granules utilize so many p97 adaptors? UFD1-NPL4 requirement is easily appreciated as it is necessary for unfolding of the ubiquitin that drives eventual unfolding of the substrate through the central pore (Cooney et al., 2019; Twomey et al., 2019). Recent studies suggest that p97 adaptors that contain UBA domains such as UBXD8 or p47 may decrease the ubiquitin threshold necessary for substrate unfolding by the p97-UFD1-NPL4 complex (Fujisawa, Polo Rivera, & Labib, 2022; Huo et al., 2025). Thus, the recruitment of multiple adaptors may allow for rapid disassembly of stress granules without the need for persistent ubiquitylation or the accumulation of long ubiquitin chains on stress granule proteins. Thus, additional adaptors may help allow for faster disassembly. It is also possible that each adaptor may recognize a unique set of ubiquitin modified substrates on stress granules. We and others have shown interaction of UBXD8 and p47 with ubiquitylated G3BP1 as it is the most abundant stress granule component, but other substrates likely exist that may be targeted by these adaptors uniquely. These remain to be identified and are the focus of future work.

---

**Supplementary Figure 1.**
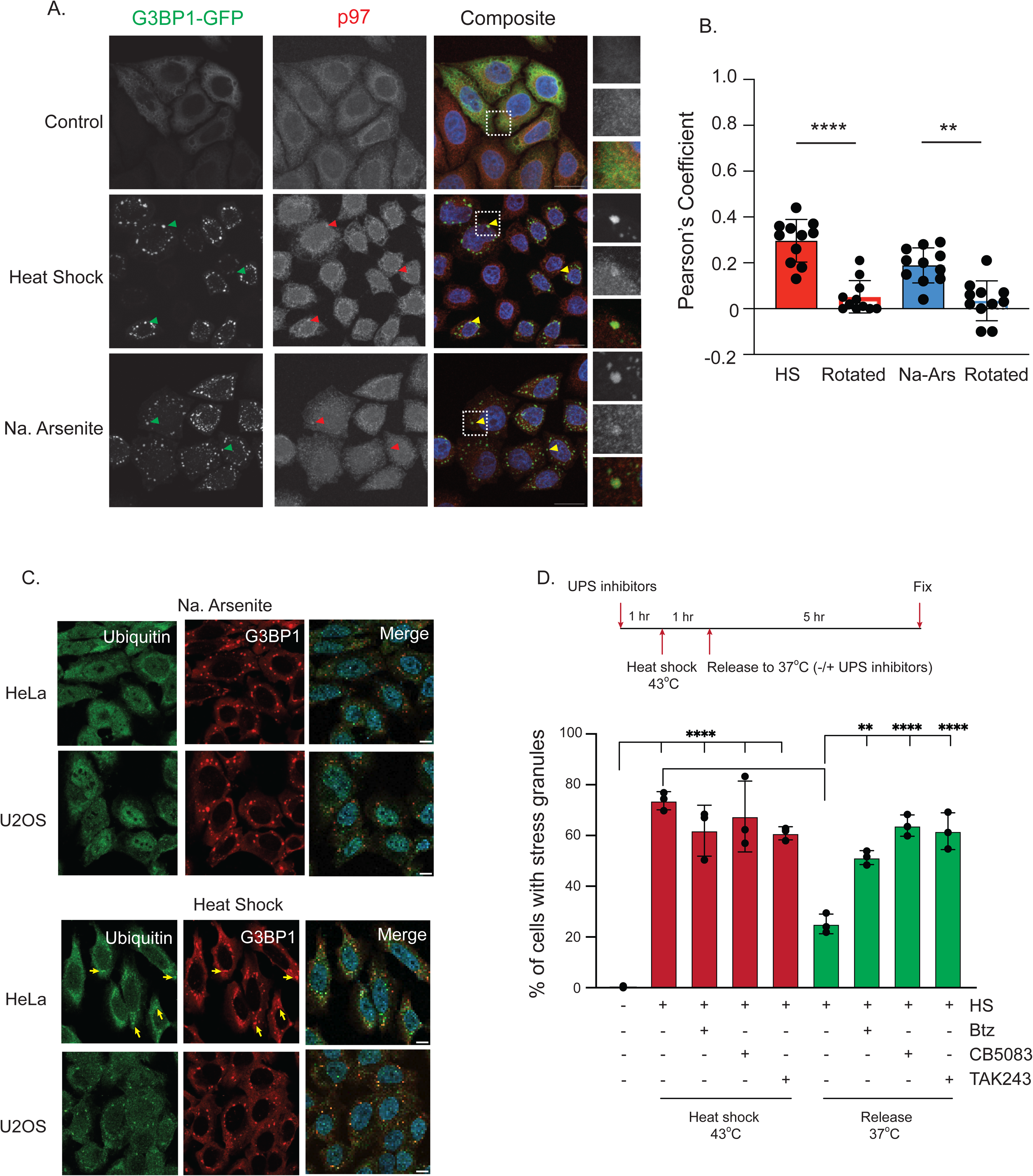
p97 is recruited to stress granules induced by heat shock and sodium arsenite. (A) Hela Kyoto cells stably expressing GFP-G3BP1 were exposed to either heat shock stress (HS, 42 °C for 1 hour) or sodium arsenite (Na Ars., 500 μM for 1 hour). Treated cells were then fixed and stained for endogenous p97 (red) which was observed to be recruited to G3BP1 positive punctate (green) following HS and Na Ars. Induced stress. Nuclei were counterstained with Hoechst. (Scale bar: 20μm) (B) Colocalization is numerically represented via the Pearson’s correlation coefficient (PCC) of p97 with G3BP1 positive punctate. The significance of correlation was determined by comparing the PCC of stress condition images to the PCC of a matched but rotated image, termed “Rotated”. Shown data was generated from 3 independent experiments where 5 images were quantified per condition. Data is represented as mean PCC +/- Standard Deviation (SD). Significance was determined via a paired t-test, p<0.05, *. (C) HeLa or U2OS cells were exposed to either sodium arsenite (500 mM for 1 hour) or heat shock (42 °C for 1 hour). Exposed cells were then fixed and stained for endogenous G3BP1 (red) and ubiquitin (green) which was observed to be recruited to G3BP1 positive punctate only following following heat shock. Nuclei were counterstained with Hoechst. (Scale bar:10μm). (D) Hela Kyoto GFP G3BP1 cells were pre-treated for 1 hour with a proteosome inhibitor (Bortezomib, Btz), p97 inhibitor (CB5083), or ubiquitin E1 inhibitor (TAK243) then exposed to 1 hour of heat shock treatment at 42 °C. A subset of cells were allowed to recover from heat shock treatment for 5 hours in the absence or presence of the above inhibitors before cells were fixed and prepared for immunofluorescent staining. Cells were imaged and the quantification of percentage of cells containing stress granules is shown. 250-300 cells were analyzed in N=3 independent experiments. Data is presented as mean +/- Standard Deviation. *,**,***,**** : P< 0.05,0.01,0.001,0.0001. Significance was determined via One-way ANOVA with Tukey’s post-hoc test. Scale bar is 10 μM.

**Supplementary Figure 2.**
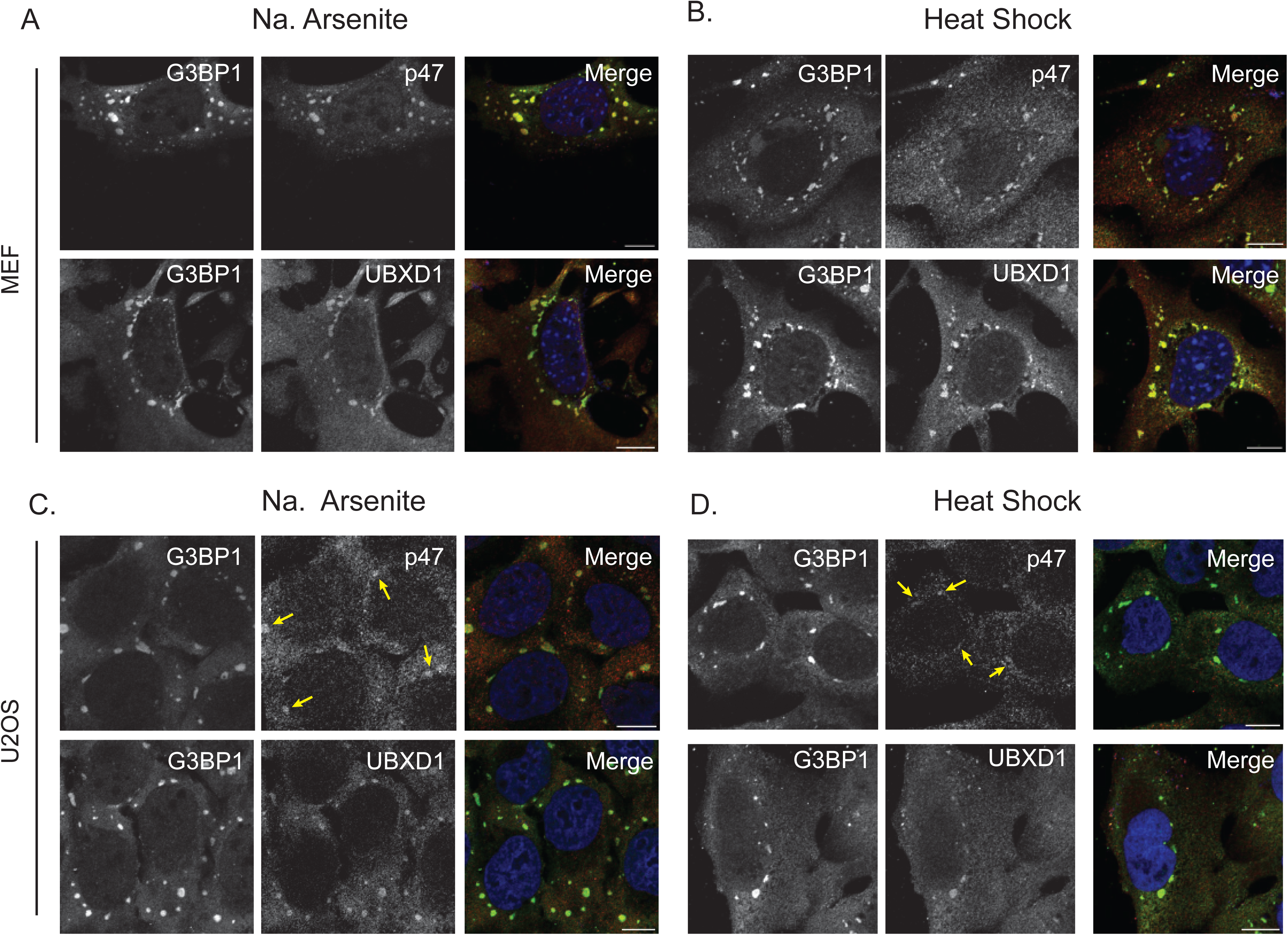
p47 and UBXD1 are recruited to stress granules in multiple cell types. (A), (B) Mouse embryonic fibroblasts (MEF) or (C), (D) U2OS cells were treated with sodium arsenite (500 μM for 1 hour) or heat shock (42 °C for 1 hour) respectively. Cells were fixed and stained for endogenous p47, UBXD1 or G3BP1. p47 recruitment to stress granules in U2Os cells is less robust than in other cell types and is indicated with yellow arrows for clarity. Nuclei were counterstained with Hoechst. N= 3 biological replicates. Scale bar is 10 μM.

**Supplementary Figure 3.**
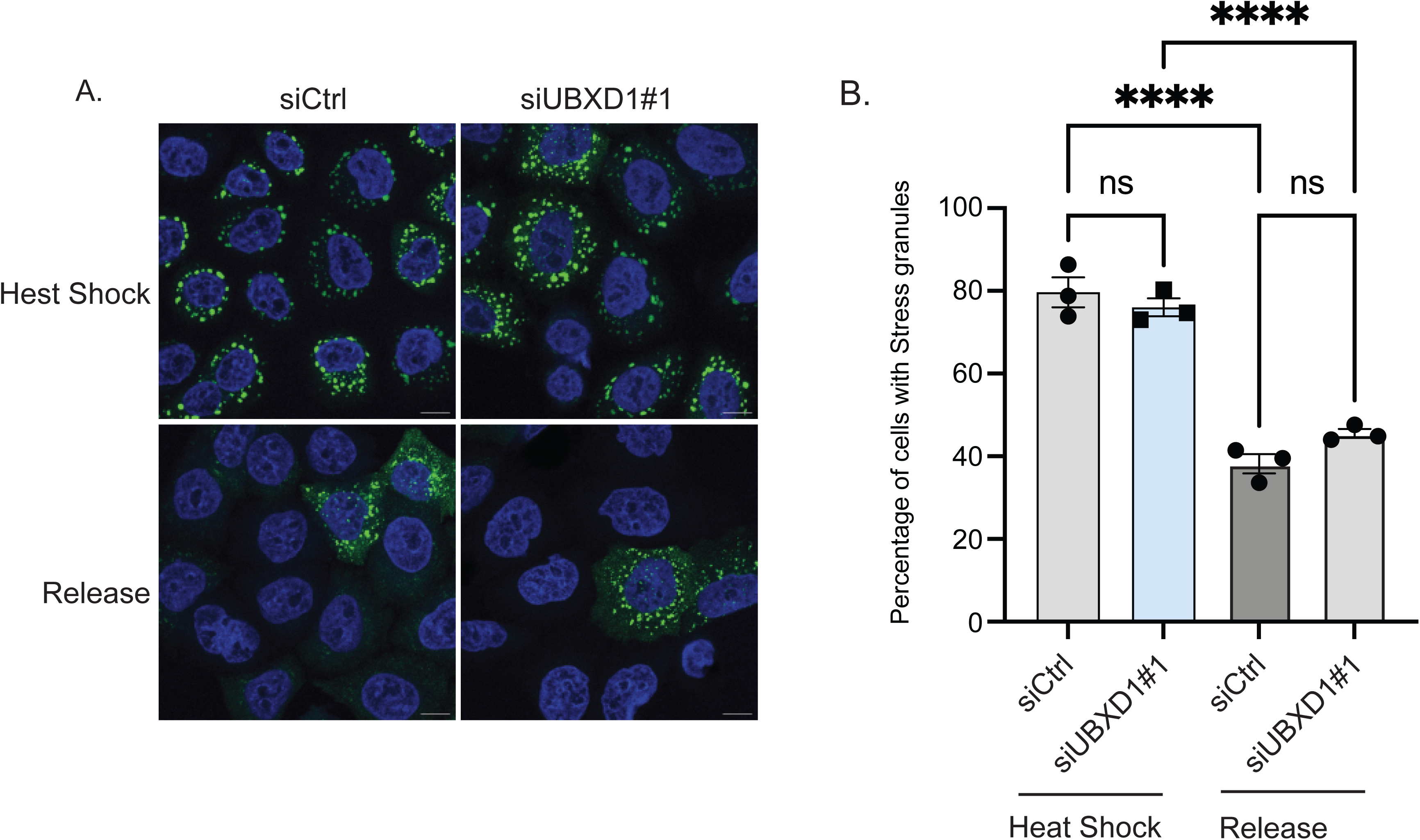
UBXD1 is not required for disassembly of stress granules after heat shock. (A) HeLa Kyoto GFP-G3BP1 cells were transiently transfected with siRNAs to non-targeting control (Ctrl) or UBXD1 for 48 hours. Immunoblot of knockdown efficiency is shown in main Figure 2A. Cells were treated with Heat Shock (42°C for 1 hour) and then released into 37°C for 5 hours. Cells were fixed and imaged for GFP-G3BP1. Nuclei were counterstained with Hoechst. (B). Quantification of number of cells with stress granules from (A). UBXD1 is not required for the dissolution of stress granules. 175-250 cells were analyzed in N=3 independent experiments. Bar graph shows mean and standard deviation. ****: p< 0.0001 as determined by One way ANOVA with Dunnett’s multiple comparisons test. ns: not significant. Scale bar is 10 μM.

## Material and Methods

### Cell Culture and Treatments

HeLa Kyoto cells were a gift from Ron Kopito Stanford. Mouse embryonic fibroblasts and U2OS (ATCC) cells were cultured in Dulbecco’s modified Eagle’s medium, supplemented with 10% fetal bovine serum (FBS) and 100 units/ml penicillin and streptomycin. Cells were maintained in a humidified, 5% CO2 atmosphere at 37 °C. HeLa GFP-G3BP1 stable cells were generated by lentiviral transduction with pHAGE-GFP-G3BP1 and lentiviral helper vectors. Stable cells were selected using blasticidin selection. Cells expressing low levels of GFP-G3BP1 (lower quartile of the total GFP signal) were collected by flow sorting.

Stress granules were induced by treating cells with heat shock (42 °C) for the indicated times (see figure legends) or with 500 μM sodium arsenite. Cells were released from heat shock by returning cells to 37 °C or from sodium arsenite treatment by washing three times in pre-warmed media and returning to drug free media for the indicated times (see figure legends). Cells were treated with 1 μM Bortezomib, 5 μM CB-5083, or 2 μM TAK-243 in the indicated experiments.

For siRNA transfections, cells were either forward, or reverse transfected with 20 nM siRNA using Lipofectamine RNAiMax (Invitrogen) in a 12- or 6-well plates according to the manufacturer’s protocol. After 24 or 48 hours, depending on the study, cells were split into 12-well plates for further analysis. 48 or 72 hours post-transfection, cells were harvested for immunoblot or fixation for immunofluorescence.

For p47 siRNA rescue experiments cells were first reverse transfected with p47 siRNA #4 and then forward transfected 24 hours later with 500 ng FLAG-p47 wildtype siRNA resistant construct for a further 24 hours using Lipofectamine 2000 (Invitrogen).

### Antibodies and Chemicals

The p97 (10736-1-AP; IF: 1:100), UBXD8 (16251-1-AP; WB: 1:2000), UBXD1 (147061-1-AP, WB: 1:1000, IF: 1:100), G3BP1 (13057-1-AP, 1:1000, IF: 1:100), NPL4 (116381-1-AP, 1:1000, IF: 1:100), antibodies were from Proteintech Inc. UBXD7 (PA561972, IF: 1:100) was from Thermo Fisher. The pan-ubiquitin (P4D1; sc8017; WB: 1:2000), β-Actin (AC-15; sc69879; WB: 1:2000), and GAPDH (O411; sc47724; WB: 1:2000) antibodies were obtained from Santa Cruz Biotechnologies. p97 (A300-589A; WB: 1:2000) was from Bethyl Laboratories. anti-FLAG (M2; F3165) was from Sigma Aldrich; WB: 1:5000. HRP-conjugated anti-rabbit (W401B; WB: 1:10,000) and anti-mouse (W402B; WB: 1:10,000) secondary antibodies were from Promega. Goat anti-Mouse IgG (H + L) Cross-Adsorbed Secondary Antibody, Alexa Fluor™ 568 (Catalog # A-11004; IF: 1:10,000), and Goat anti-Mouse IgG (H + L) Cross-Adsorbed Secondary Antibody, Alexa Fluor™ 488 (Catalog # A-11001; IF: 1:10,000) were purchased from Thermofisher Scientific. Bortezomib (72-825) and CB-5083 (73-795) were from Tocris, TAK-243 (S833411) from Selleckchem.

siRNAs used in this study: p47 siRNA #1 (D01700001) and #4 (D01700004) and UBXD1 siRNA #1 (D008785) were from Dharmacon, UBXD8 siRNA #0 (s23260) was from Ambion (Thermo Fisher Scientific). siControl (siC001) was from Millipore Sigma. FLAG-p47 was cloned by Gateway cloning from the Orfeome collection.

### Immunofluorescence and Microscopy

Cells were plated on # 1.5 glass coverslips in a 12-well plate. Following indicated treatments, cells were fixed in 4 % paraformaldehyde (PFA) (15710-S Electron Microscopy Sciences) diluted in PBS for 15 minutes at room temperature. Next, cells were washed in PBS and permeabilized in ice-cold 100% methanol at − 20 °C for 10 min. Cells were washed three times in PBS and incubated in a blocking buffer (1 % BSA, 0.3 % Triton-X100 for 1 hour at room temperature. Primary antibodies were diluted to the indicated concentrations in blocking buffer, and coverslips were incubated overnight at 4 °C in a humidified chamber. Coverslips were washed three times in PBS and incubated in secondary antibodies diluted to the indicated concentrations in a blocking buffer for 1 hour at room temperature. The secondary antibody solution was replaced with Hoechst diluted in blocking buffer and incubated for 5 minutes at room temperature. Coverslips were washed three times with PBS and mounted to slides with ProLong Gold antifade mounting media (P36930 Invitrogen). All images were collected using a Zeiss LSM800 confocal microscope equipped with Airyscan using Zeiss Zen Black 2.3 software. Images were taken with optimal pinhole a 63× oil immersion objective (NA 1.4) at 25 °C. The indicated fluorophores were excited with a 405, 488, or 594 nm laser line.

### Image analysis

Images were analyzed using FIJI (https://imagej.net/fiji). Stress granule number per cell and size were measured using an automated image analysis script (Aggrecount), allowing segmentation and single-cell resolution (Klickstein, Mukkavalli, & Raman, 2020). For colocalization, briefly, images were divided into three channels: (1) p97/adapter channel for p97, p47, UBXD8, UBXD1, or NPL4 (2) G3BP1 and (3) Hoechst. Z stack images for each channel were collapsed via Max Intensity Projection. A background subtraction was performed for each channel with the rolling ball algorithm. To create a mask of the boundaries of the stress granules punctate, gentle blur and thresholding were performed on the G3BP1 channel. Lastly, the Fiji plugin for colocalization analysis, Coloc2, was used to determine the Pearson’s correlation coefficient (PCC) between the p97/adapter and G3BP1 channel. To determine if the magnitude of PCC returned was due to random pixel colocalization, we also generated the PCC from a paired set of images where the channel for the G3BP1 channel was rotated 90°. A new mask was created from this 90° rotated imaged (“rotated”) and used to generate the PCC against p97/adapter and G3BP1.

### Immunoprecipitation, SDS-PAGE analysis and Western blotting

To evaluate knockdowns, cells were lysed in mammalian cell lysis buffer (50 mM Tris-Cl, pH 6.8, 150 mM NaCl, 0.5% Nonidet P-40, HALT Protease inhibitors (Pierce), and 1 mM DTT). Cells were incubated at 4 °C for 10 min and then centrifuged at 19,000 × g for 15 min at 4 °C. The supernatant was collected, and protein concentration was estimated using the DC protein assay kit (Biorad).

For immunoprecipitation experiments, cells were lysed as in stress granule lysis buffer (Gwon et al., 2021) (25 mM Tris-HCl, pH 7.4, 150 mM NaCl, 1% NP-40, 1 mM EDTA, 5% glycerol; 20 mM *N*-ethylmaleimide (NEM) (Sigma-Aldrich E3876), 50 μM PR-619 (Sigma-Aldrich, 662141) and proteinase inhibitor cocktail (Roche 1183617001). Lysates were centrifuged at 4°C for 15 min at 20,000*g*. Total protein was estimated by BCA assay (Biorad). Equal amounts of total protein were used for immunoprecipitation with the indicated antibody along with 30 μl of a 1:1 slurry of protein G Sepharose beads (Pierce) with end to end rotating overnight. Lysates were washed 5 times with 1 ml of stress granule lysis buffer with 300mM NaCl, resuspended in 30 μl of 2X Laemmli buffer, boiled at 95 °C and resolved on SDS PAGE.

For immunoblotting of lysates: The protein concentrations were estimated by the BCA assay (ThermoFisher Scientific 23225). Samples were loaded into 10 % SDS-PAGE gel and transferred into PVDF membrane. The membranes were blocked in 5 % milk in TBS-0.2% Tween20 and incubated with the primary antibody in blocking solution overnight at 4 °C. The membrane was washed three times with TBS–0.2% Tween20 and incubated with respective secondary antibodies for 1 h. Membranes were rinsed three times with TBS–0.2% Tween20 and developed with Bio-Rad Western ECL Substrate (1705061) in the dark and imaged using Bio-Rad Chemi-Doc Imaging system (Bio-Rad). Images were analyzed using ImageJ 1.52q Analyze gels tool.

### Statistics and reproducibility

For all experiments, n ≥ 3 or more biological replicates for each condition examined. Fold changes, SEM, SD, and statistical analyses were performed using GraphPad Prism version 9.4.1 for Windows (GraphPad Software). Statistical tests and N values are mentioned in the figure legends.

## Acknowledgments

This work is supported by the R01 GM127557, R21 NS123631 and Muscular Dystrophy Association grants to M.R., and K12GM133314 for the Tufts IRACDA Program to M.J.

## Respective Contributions

M.A.J, R.K, S.M and M.R conceived the study, M.A.J, R.K, S.M, L.N and M.R performed the experiments and M.A.J, R.K, S.M and M.R analyzed the data. MR wrote the manuscript with input from M.A.J, R.K, S.M.

## Request for reagents

Please contact the corresponding author, M.R for reagent requests.

